# MicroRNA-sensing plasmid system for dynamic control of functional ion channel expression

**DOI:** 10.64898/2026.02.04.703753

**Authors:** Joschua Geuter, Naxi Tian, Jordan Brown, Stephanie Schorge, Gareth Morris

## Abstract

Gene therapy offers the potential for long-term treatment or cures for a range of chronic diseases. However, permanent gene therapy expression may not be desirable. Efforts have been made to create systems which can be switched on/off by stimuli including light, designer drugs, or cellular contexts such as increased electrical activity. Here, we designed a novel plasmid system in which ion channel expression, and therefore function, is regulated by microRNA (miR) – an endogenous class of short noncoding RNAs which negatively regulate gene expression via binding the 3’ untranslated region of target transcripts. We modified an existing voltage-gated potassium channel gene therapy with a binding cassette for miR-193a-3p, and transfected this ‘miR-193-OFF’ system in neuro2A cells. Co-transfection with an inhibitor or mimic of miR-193a-3p respectively enhanced or repressed expression of our transgene, assessed using a GFP marker. Using whole-cell voltage clamp, we observed enhanced voltage-gated potassium currents in cells co-transfected with a miR-193a-3p inhibitor, compared with a non-targeting control oligonucleotide. Together, this demonstrates the concept of a novel miR-mediated molecular switch which can bias therapeutic ion channel expression based on a specific miR signal. As miRs are a ubiquitous molecular mechanism, our approach could be applied to a wide range of cellular and disease contexts, potentially expanding gene therapy to new patient populations.

## Introduction

Gene therapy offers the prospect of long-lasting therapeutic intervention for patients with otherwise treatment-resistant chronic neurological diseases^1^. In brief, gene therapy uses viral vectors, typically adeno-associated virus (AAV) or lentivirus (LV), to deliver therapeutic DNA plasmids into target cells. In most experimental settings, gene therapies are injected directly into a target area in the brain and do not spread far from their injection site, with AAVs having a wider spatial distribution - but smaller packaging capacity^2^ - than LVs due to their smaller physical size. Certain viral serotypes confer preferential transduction of different cell types, and promoter and enhancer elements can be incorporated into gene therapy plasmids to further bias transgene expression to specific cell populations. These properties offer a certain level of control over where in the brain, and in which cells, gene therapies are expressed. ‘Classical’ gene therapy strategies, therefore, permit very long-term constitutive expression of therapeutic transgenes to treat chronic neurological diseases. Indeed, gene therapy treatments for rare monogenetic disorders are already clinically available (such as Zolgensma for spinal muscular atrophy^3^), and promising experimental data exists to support the use of gene therapies in various brain conditions, including epilepsy^1–4–5^, Parkinson’s disease^6^, and chronic pain^7^.

However, permanent constitutive gene therapies may also raise important safety concerns: the therapy may not need to be permanently activated, and any adverse effects caused by unnecessary activation would be irreversible in most scenarios (with the possible exception of epilepsies eligible for surgical resection that could physically remove the transduced brain region). There is a critical unmet need for dynamic gene therapy tools that can modify their expression in response to cellular or exogenous stimuli. Experimental approaches include optogenetics^8^, in which light is delivered into the brain to activate light-sensitive ion channels, and chemogenetics^9–10^, in which therapeutic ion channels are activated by a specific ligand which must be administered via medication. In both of these strategies the treatment is only active in the presence of an external stimulus, which can be removed if safety concerns arise. However, such approaches do not respond to endogenous changes to cellular environments. A recent advance used a promoter sequence from the immediate-early gene *c-Fos* in order to develop an ‘activity-dependent’ gene therapy^11^. This has substantial therapeutic potential in diseases such as epilepsy, in which changes in brain activity are a hallmark of the condition, and therefore the therapy responds on-demand to the dynamics of the disease itself. Novel strategies, responding to other changes in cellular context during brain disease, would represent substantial breakthroughs in the development of disease-driven on-demand gene therapies, especially those where cellular excitability is not a hallmark of disease.

One such cellular context which could be exploited to regulate gene therapy expression is microRNA (miR) expression. miRs are ∼22 nucleotide noncoding RNAs that negatively regulate gene expression through binding complementary sites in the 3’ untranslated region (UTR) of target mRNAs^12-14^. Changing miR levels are closely associated with various brain diseases and in some cases are considered to be biomarkers and/or therapeutic targets ^15–17^. Taken together, miRs offer a natural mechanism to modify gene expression, and the expression of miRs themselves is tightly linked to disease contexts. To generate a proof-of-concept, we modified the published ‘engineered potassium channel^5^’ (EKC) epilepsy gene therapy, cloning a binding cassette for miR-193a-3p^18^ into the 3’ UTR. By incorporating these miR binding sites, we created a system where high miR-193a-3p levels repress ion channel expression and function through miR-mediated silencing, while reduced miR-193a-3p levels allow enhanced therapeutic gene expression – effectively creating a miR-responsive molecular switch. Using a cell-based assay, we demonstrate specific and bidirectional changes in EKC expression and channel function in response to changing miR-193a-3p expression.

## Results

### miR-193-3p sensitive plasmid design

We based our approach on the previously successful ‘engineered potassium channel’ (EKC) plasmid^5^ and inserted a binding cassette for miR-193a-3p into the 3’ UTR, directly following the stop codon (Figure 1A,B). The synthetic binding cassette consisted of three consecutive repeats of the sequence TGTAGGCCAGTT, based on a naturally occurring miR-193a-3p binding region in the 3’ UTR of the *PTEN* gene^18^. The sequence TGTAGGCCAGTT provides full complementarity, and auxiliary binding, to miR-193-3p’s seed region. Given the natural ability of miRs to repress their target genes, our plasmid (‘miR-193-OFF’) was designed to respond dynamically to changes in miR-193a-3p expression: when miR-193a-3p is low and does not repress the plasmid, the therapy is more active; when miR-193a-3p is high, it represses the plasmid and the therapy is reduced.

**Figure 1:**
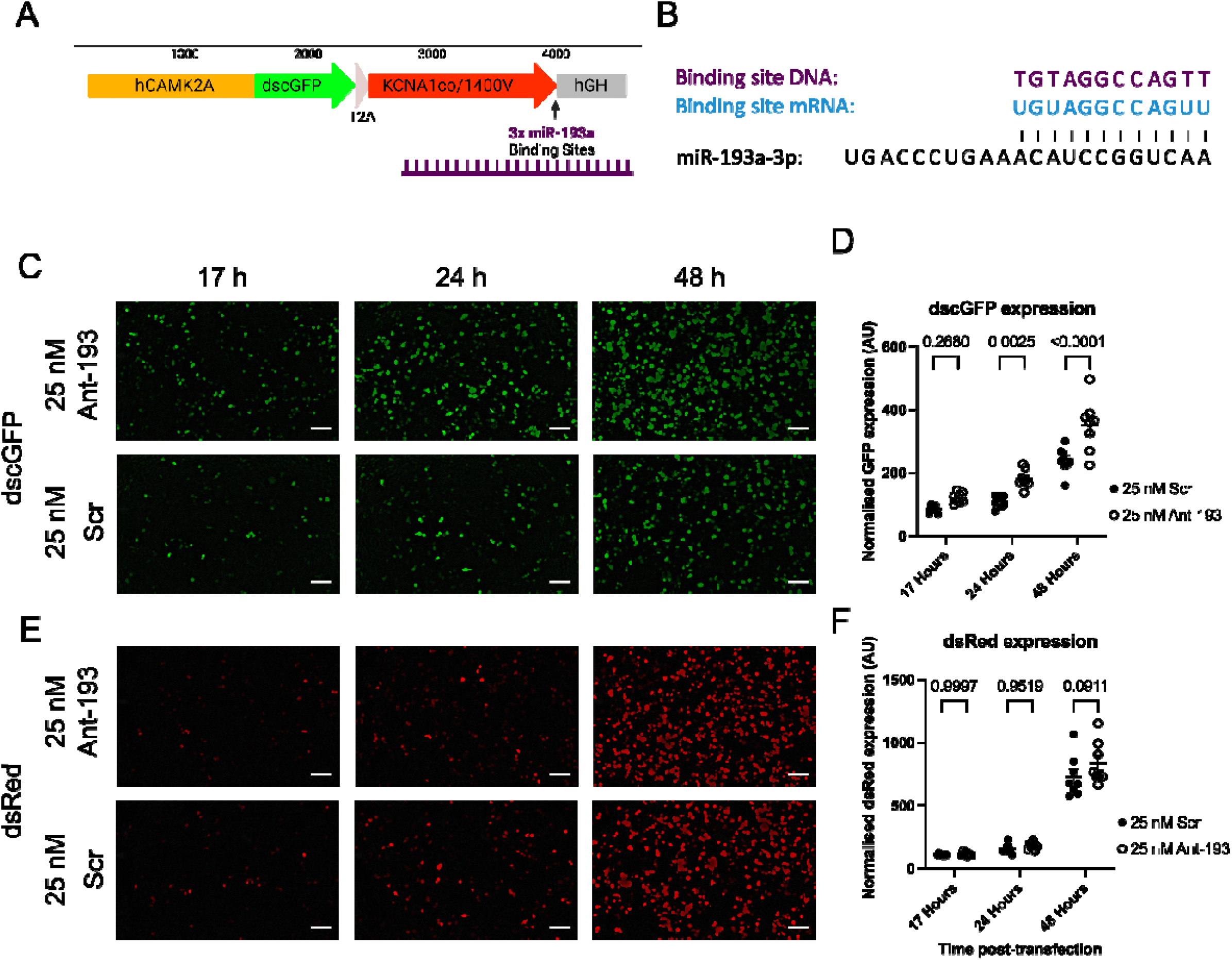
Ant-193a-3p mediates specific de-repression of miR-193-OFF plasmid. **A–** DNA plasmid design for miR-193-OFF. Design is based on a published successful gene therapy^5^ and modified to include a synthetic miR-193a-3p binding cassette^18^. **B**–Sequences of synthetic binding cassette DNA and RNA, indicating complementarity to miR-193a-3p. **C** – Neuro2a cells co-transfected with miR-193-OFF (with dscGFP marker), dsRed (non-miR-sensitive control) and 25 nM of either ant-193a-3p (‘ant-193’) or a scrambled non-targeting antimiR control (‘Scr’). Representative live cell fluorescence images acquired at different timepoints post-transfection show enhanced dscGFP expression in ant-193 treated cells, relative to control. Scale bar: 100 µm) **D** – Statistical analysis shows that a significant differential expression of dscGFP emerges at 24 hours post-transfection and is enhanced at 48 hours (RM two-way ANOVA with Sidak’s multiple comparisons; p values marked on graph). **E** – dsRed live cell images, taken from the same samples and fields of view as the dscGFP images in C. **F** – Statistical analysis indicates that dsRed, which is not sensitive to miR-193a-3p, does not exhibit significantly different expression at any time point (RM two-way ANOVA with Sidak’s multiple comparisons; p values marked on graph).

### Dose-dependent bi-directional changes in plasmid expression with changing miR-193-3p expression

To demonstrate the ability of our plasmid system to respond to changing miR-193a-3p expression, we used cultured neuro2A cells co-transfected with miR and either a locked nucleic acid antimiR (ant-193; inhibiting miR-193a-3p function), a non-targeting scramble control (Scr), or a synthetic miR-193a-3p mimic (overexpressing miR-193a-3p). All transfections further contained a miR-193a-3p-insensitive dsRed fluorescent plasmid as a control to demonstrate the specific effects of altered miR-193a-3p on miR-193-OFF. In the first instance, we used fluorescence imaging of the destabilised copGFP (dscGFP) marker (Figure 1C) as a semi-quantitative readout of plasmid expression. In each experiment, we imaged eight separate random regions of interest for each well at 17, 24 and 48 hours post-transfection, measured the fluorescence intensity for each ROI, and averaged these eight values to give an average fluorescence intensity value for each well (with each well as a biological replicate). In control conditions (miR-193-OFF transfected without ant-193, Scr, or mimic), dscGFP expression increased over time as expected, and miR-193-OFF did not cause toxicity, indicating broad safety. For cells co-transfected with 25 nM ant-193, dscGFP expression was significantly enhanced relative to Scr control within 24 hours post-transfection (Figure 1C,D). The magnitude of differential expression increased at later timepoints. Importantly, dsRed fluorescence was unchanged between ant-193 and Scr treatments (Figure 1E,F), highlighting that the changes in gene expression observed were specific to our miR-193a-3p-sensitive plasmid design and not due to general changes in cellular proliferation or protein translation.

To confirm the bi-directional dynamics of our system, we then used varying concentrations of a miR-193a-3p mimic (mim-193), seeking to repress miR-193-OFF expression through mim-193 binding to the plasmid (Figure 2). Using the same strategy as above, we demonstrated a largely dose-dependent repression of dscGFP signal by mim-193, which emerged at 48 hours posttransfection and was more prominent after 48 hours. Again, miR-193a-3p-insensitive dsRed was not repressed by the mimic, showing consistent expression across all concentrations of mim-193 within each timepoint. Together, Figures 1 and 2 show that our miR-193-OFF system mediates bidirectional changes in transgene expression, following corresponding changes in miR-193a-3p levels.

**Figure 2:**
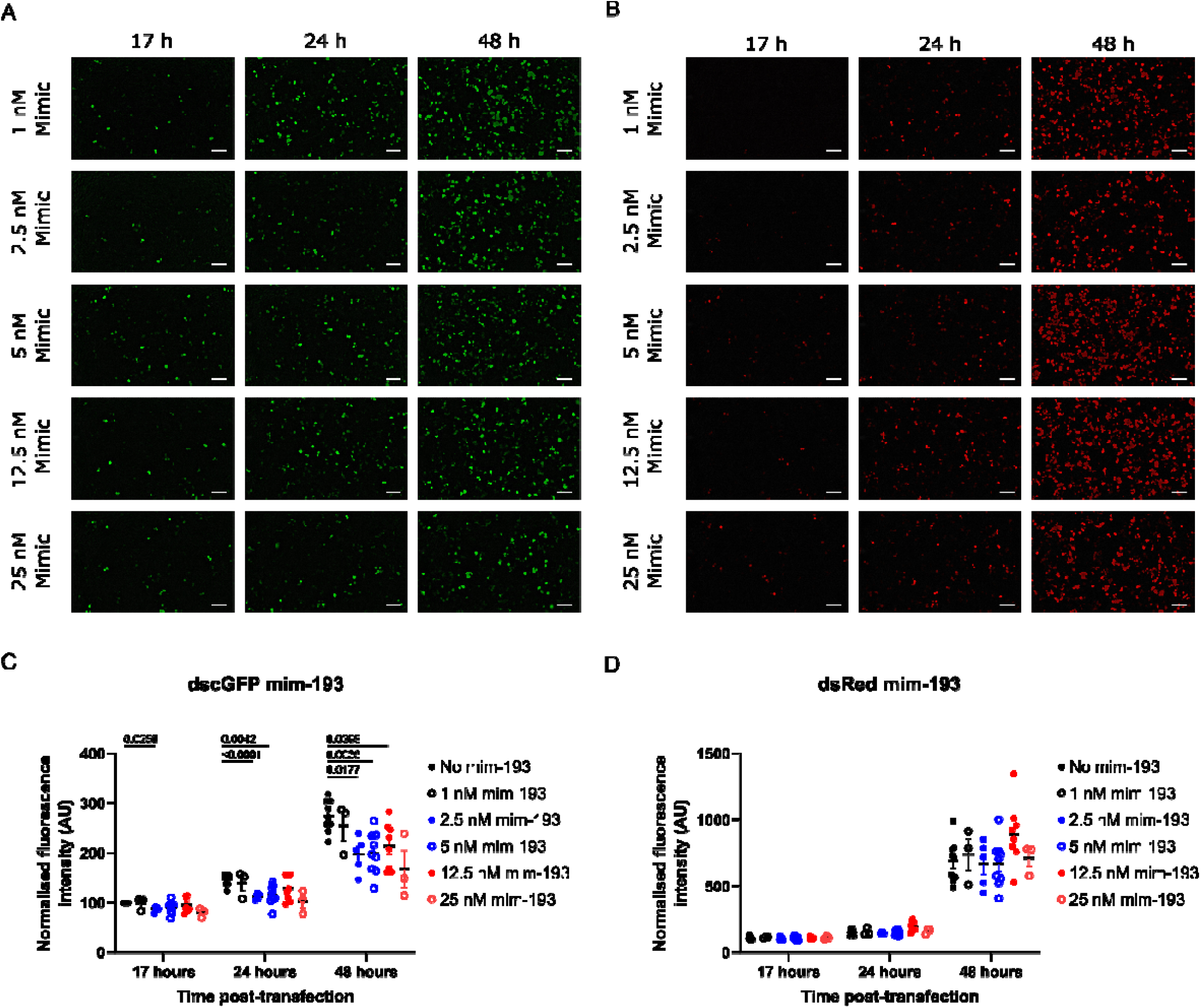
Specific and dose-dependent repression of miR-193-OFF expression by mim-193. Neuro2A cells were co-transfected with miR-193-OFF (dscGFP reporter), dsRed (non-miR-sensitive control) and varying concentrations of mim-193. **A** – miR-193-OFF expression, measured using dscGFP expression, decreased in a dose-dependent manner as mim-193 concentration increased. Scale bar: 100 µm **B** – dsRed expression was not altered by mim-193, indicating a specific response of the miR-sensitive plasmid to mim-193. Scale bar: 100 µm **C** – population fluorescence data (normalised to ‘no mim-193 17 hours’ value) for dscGFP demonstrates largely dose-dependent repression of miR-193-OFF by mim-193 (mixed two-way ANOVA with Dunnett’s multiple comparisons; significant p values at α = 0.05 marked on graph). **D** – In contrast, dsRed (which does not have a binding domain for miR-193a-3p) is insensitive to mim-193 at all timepoints (mixed two way ANOVA with Dunnett’s multiple comparisons; no p values were significant at α = 0.05).

### Modifiable voltage-gated potassium channel functional expression in response to miR-193-3p inhibition

Having explored the kinetics of miR-193-OFF using a dscGFP reporter readout, we next verified that we could modify functional ion channel expression using this system, thereby demonstrating that the therapeutic EKC mechanism can be modulated by a miR. Using whole-cell voltage clamp, we first ensured that we could record robust EKC-mediated currents (I_ekc)_ from neuro2A cells transfected with miR-193-OFF (identified by the presence of dscGFP signal), in the absence of miR inhibitors (Figure 3A-C). For completion, we also confirmed that non-transfected cells (from the same coverslips, but with no observable dscGFP fluorescence) did not exhibit voltage-sensitive potassium currents. As expected, transfected cells showed significantly higher current density than non-transfected cells (Figure 3B), and only transfected cells showed a non-linear I-V curve (Figure 3C), indicative of voltage-sensitive channel gating. Finally, we assessed EKC current density in cells which were co-transfected with miR-193-OFF and either 25 nM ant-193 or 25 nM Scr (Figure 3D-F). Whilst both groups of cells were able to generate voltage-gated potassium currents (Figure 3D), cells treated with ant-193 exhibited significantly larger current densities than Scr controls (Figure 3E,F), indicating that functional EKC expression and currents can be controlled via miR levels. This finding shows that we can create dynamic synthetic gene delivery systems by incorporating miR binding cassettes - designed to respond to particular cellular contexts - into the 3’UTR.

**Figure 3:**
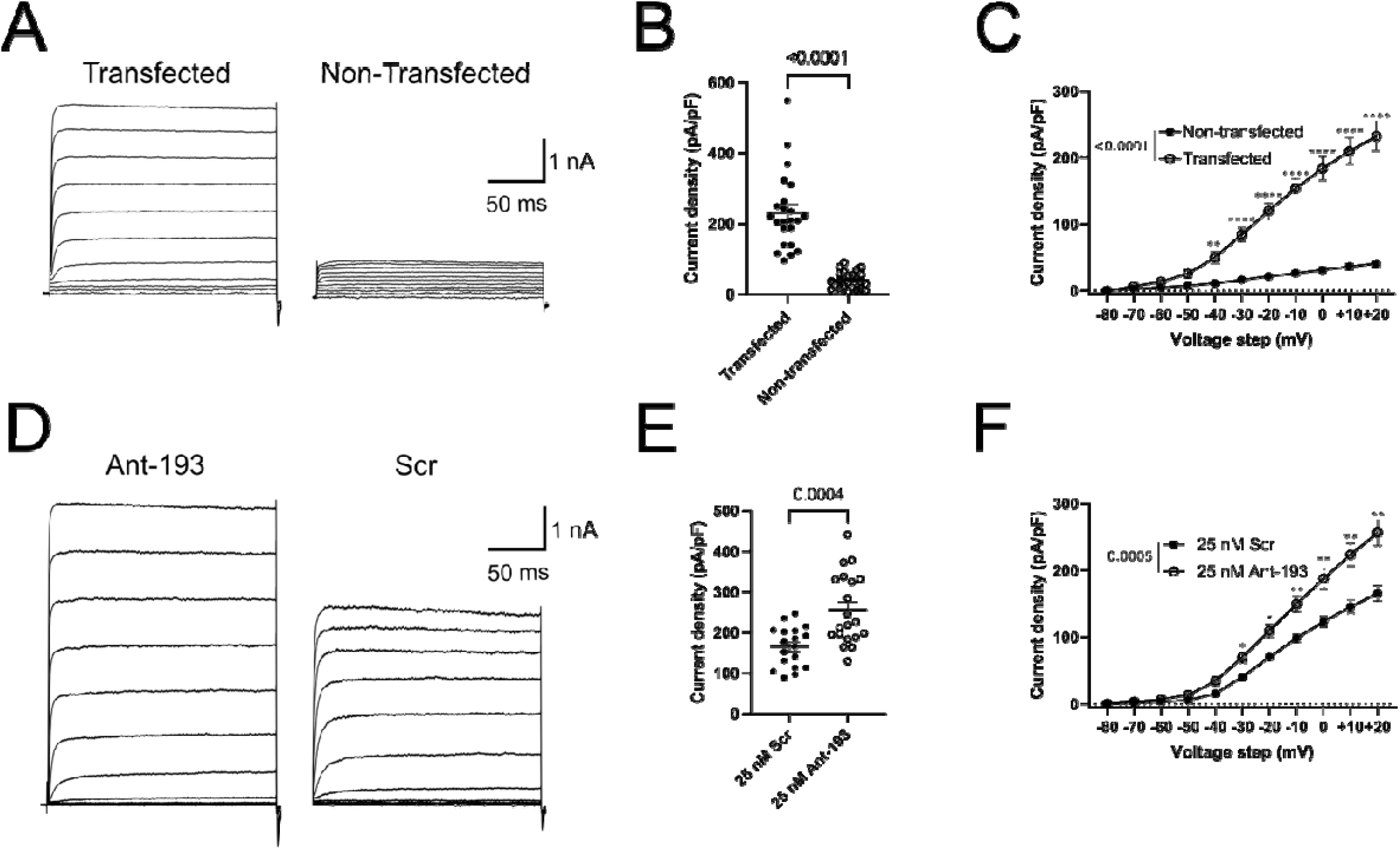
Enhanced voltage-gated potassium channel function in response to miR-193a-3p inhibition. Neuro2A cells were transfected with 25 nM ant-193 (or Scr control) and used for voltage-clamp recordings of V-gated potassium channel currents at 24-30 hours post-transfection. **A-C**: cells were transfected with miR-193-OFF only. **A**: Only transfected cells (identified using dscGFP label) exhibited voltage-dependent K^+^currents, indicated by representative traces from both transfected and non-transfected cells. **B** – Fluorescent cells had significantly greater current density than non-fluorescent cells (scatterplot shows responses to +20 mV voltage step; Student’s t-test). Using the same V-clamp protocol in non-transfected cells, we confirmed that there was no native Kvfunction in our cell line, shown by a linear I-V relationship (**C** – mixed ANOVA with Sidak’s multiple comparisons; p values marked on graph) that reflects ohmic resistance of the cell membrane only. **D-F:** in a separate experiment, cells were co-transfected with miR-193-OFF and either ant-193 or Scr control. In each group, we selected transfected (dscGFP-positive) cells for recordings. **D** – Representative voltage-gated K^+^currents in cells co-transfected with ant-193 or Scr. Ant-193-treated cells showed markedly enhanced l_Kv_ (**E** – responses to +20 mV voltage step, Student’s t-test; **F** – I-V curve for all voltage steps), demonstrating that ion channel function in our plasmid is modulated by changing miR-193a-3p expression (mixed ANOVA with Śidák’s multiple comparisons; p values marked on graph).

## Discussion

Here, we demonstrate a dynamic gene therapy expression system where synthetic ion channel expression can be modulated bi-directionally in response to endogenous miR signals. As a proof-of concept, we modified an existing promising pre-clinical *Kcna1*-based epilepsy gene therapy plasmid, adding a new level of control mediated by miR-193a-3p.

### A new level of gene therapy control

Concerns remain that constitutive ‘classical’ gene therapy expression may pose safety risks since adverse effects could not be reversed in the majority of cases. As such, various gene therapy strategies have been developed which respond to certain stimuli, offering a level of control over their expression. Strategies include optogenetics and chemogenetics and, more recently, cell autonomous approaches such as ‘activity-dependent’ gene therapy^11^. The latter circumvents the need for exogenous control of the gene therapy by using an activity-driven promoter to bias gene therapy expression to the most active neurons – a strategy particularly amenable to epilepsy. In this work, we developed a novel expression system in which a gene therapy can be modulated by changing levels of an endogenous miR. This has several advantages. First, it does not require exogenous activation (unlike opto- and chemo-genetics), removing the need for invasive delivery of light to potentially deep brain structures (optogenetics), or administration of a designer ligand (chemogenetics). Second, miR-regulation of gene expression is ubiquitous to all cells and, therefore, our strategy may have utility in a broad range of tissues, compared with an activity-dependent strategy which exploits enhanced activity in excitable cells only. Third, there are many hundreds of different miRs in mouse and thousands in human. Thus, our initial proof-of-concept could be reconfigured to respond to changes in specific individual miRs, or combinations of them. Strategic selection of a miR biomarker signature may be highly context- or disease-specific.

### A novel system to modulate ion channel expression

Certain miRs are known to have physiological roles in fine-tuning the expression of genes encoding for ion channels^19^. For example, miR-324-5p represses *Kcnd2*^20^– encoding K_v_4.2 – and miR-335-5p targets multiple voltage-gated sodium channel genes^21^. Therefore, it is known that miRs form part of a natural homeostatic mechanism to maintain neuronal excitability within a physiological range^19,22^. Here, we exploit this natural function of miR to add a layer of control to the expression of an exogenous ion channel plasmid. Previous studies have used a similar approach as a semi-quantitative readout of miR expression by cloning combinations of miR binding sites into the 3’ UTR of luciferase reporter gene ^23,24^. This strategy permitted the authors to study the temporal kinetics of specific miRs *in vivo*. However, to our knowledge, our study is the first to demonstrate a dynamic exogenous ion channel expression system using a miR-based switching mechanism. The system is reconfigurable with different promoters, transgenes, and miR-binding cassettes and is therefore amenable to a wide range of applications where expression of a particular miR is a proxy of a cellular state in which expression is desirable.

### Limitations

Whilst our system is able to mediate dynamic transgene gene expression, the system was not fully switched off even in the presence of relatively high mim-193 concentrations (Figure 2A,C). This is consistent with the physiological roles of miRs, which fine-tune gene expression rather than acting as binary ON/OFF switches. However, in certain applications, it may be preferable to fully switch off transgene expression when it is not required. This is beyond the functionality of our proof-of-concept, but it is conceivable that miR-mediated transgene repression could be enhanced by different configurations of binding sites.

Although not intended as a primary mechanism of our system, miR-193-OFF may lead to ‘sponging’ of the miR which is used to trigger the switching mechanism. Briefly, miRs typically mediate target repression via one of two canonical mechanisms – mRNA degradation, or steric blocking. In the case of steric blocking, miRs may form stable duplexes with their mRNA targets, therefore ‘sponging’ the miR and preventing it from engaging with its physiological targets. Due to the deliberate perfect complementarity between miR-193a-3p and our synthetic binding cassette, we anticipate that miR-193-OFF should favour the degradation mechanism, thereby minimising sponging effects. However, further study is required to interrogate the exact mechanism of gene repression by miR in our system.

## Conclusions

*We* have demonstrated proof-of-concept for a dynamic and bidirectional miR-modulated transgene expression system for ion channels. This raises the prospect of gene therapy expression which is highly specific to certain cellular or disease contexts, alleviating potential safety concerns about the permanent constitutive expression of gene therapy plasmids.

## Materials & Methods

### miR-193-OFF plasmid development

*We* based our approach on the EKC plasmid, developed in Stephanie Schorge’s lab^5^, which expresses a modified and codon-optimised version of the voltage-gated potassium channel K_v_1.1. In this work, EKC was expressed within an AAV2 plasmid backbone, containing inverted terminal repeats (ITRs), and under the control of a human CAMK2A promoter. For optical readout of transgene expression, destabilised dscGFP was coupled to EKC via a T2A peptide.

We identified the miR-193a-3p binding motif GGCCAGU from the 3’ UTR of the human *PTEN* gene^18^, which binds the miR-193a-3p seed region 2-ACUGGCC-8. We added the auxiliary binding nucleotides to create a full 1x miR-193 binding motif: TGTAGGCCAGTT, and repeated this 3x to form a miR-193 binding cassette: TGTAGGCCAGTTTGTAGGCCAGTTTGTAGGCCAGTT. This cassette was cloned into the EKC plasmid immediately downstream of the ATG stop codon, in order to place the expression of the exogenous EKC plasmid under the influence of miR-193a-3p (‘miR-193-OFF’)-The full miR-193-OFF EKC plasmid is shown diagrammatically in Figure 1.

### Cell culture and transfection

Mouse neuro2a (N2A) cells were cultured in a humidified 5% CO2 atmosphere at 37 °C in DMEM (Gibco 41966-029) supplemented with 10% Fetal Bovine Serum (Sigma F7524), 1% nonessential amino acids (Sigma M7145-100mL), and 1% Penicillin/Streptomycin (Gibco 15140-122). Cells were split every 2–3 days to maintain logarithmic growth and kept for up to 20 passages. For splitting, cells were washed with 2 mL Hank’s Balanced Salt Solution (HBSS, Gibco 14170-112) and incubated for 5 minutes at 37° C with 0.05 % trypsin-EDTA (Gibco 25300-054). Trypsin reaction was halted using 3 mL prewarmed DMEM + FBS. After centrifugation (1 minute, 1000 rpm), cells were resuspended in 1 mL cell culture medium. Subsequently, one-eighth of the cell solution was reseeded in a fresh T75 flask containing 20 mL medium.

Cells for transfection were cultured as described and seeded in a 6-well dish with 4 mL of cell culture medium 24 h before transfection, sufficient to achieve ∼30 % total confluency during transfection, which facilitated optimal transfection efficiency for this work. Chemical transfections used 6 μL TurboFect transfection reagent (ThermoFisher, R0534) in 400 μL serum-free OptiMEM (Gibco 31985-062) per well. Plasmid DNA for miR-193-OFF and dsRED (as miR-insenstive control) were added to achieve a final concentration of 0.5 μg DNA per plasmid per well. Locked nucleic acid miR inhibitors with phosphorothioate backbone were purchased from Qiagen with the following sequences: AntimiR-193a-3p (‘Ant-193’; 5’-TGGGACTTTGTAGGCCAGT-3’), non-targeting scrambled control (‘Scr’; 5’-TAACACGTCTATACGCCCA-3’). Experiments involving miR mimic used synthetic miR-193a-3p (‘mim-193’) with the sequence 5’-AACUGGCCUACAAAGUCCCAGU-3’.

The respective oligonucleotides were added to the transfection solution to achieve a final concentration in the well of 12.5 or 25 nM for ant-193 or Scr, or 1, 2.5, 5,12.5, or 25 nM for mim-193, based on manufacturer’s guidelines. After a 20-minute incubation, the transfection solutions were added to the wells. Transfection medium was replaced the following morning, approximately 17 hours post-transfection, prior to initial imaging. All transfections were conducted blinded to the experimental conditions.

### Fluorescence imaging

Transfected cells were imaged at 17-, 24-, and 48-hours post-transfection using an Olympus IX71 microscope (10x objective) equipped with a Kiralux Camera and ThorCam software (Version 3.7.0.6). Eight individual regions of interest were imaged for each condition and time point, and values presented are an average of the eight images. Imaging sites within the well were randomly selected and ensured to contain comparable cell density across conditions. Sequential dsRED (545/583 nm excitation/emission) and dscGFP (470/508 nm excitation/emission) fluorescence images were obtained using the CoolLED fluorescence illuminator. Image acquisition was conducted blinded to the experimental conditions within each well.

### Image analysis

Fluorescence intensity in dscGFP-positive cells was quantified using FIJI^26^ (Version 1.54) by calculating integrated density, which summed pixel intensities across the image and divided by image area to present total fluorescence. Background fluorescence was subtracted by creating a Gaussian-blurred version of the fluorescence image (sigma width 30 pixels) and subtracting it from the original image prior to integrated density calculation. dsRed images were analysed using integrated density calculation, with background fluorescence derived from the minimum fluorescent intensity within each image and subtracted, using skimage (Version 0.21.0) and NumPy (Version 1.25.1) packages in Python (Version 3.9). Mean integrated density was calculated across eight replicates for each condition and timepoint. To normalize between transfections, mean integrated density values were divided by the respective 17-hour negative control value, presenting all data as a percentage of control minimal fluorescence for each trial.

### Patch clamp

Neuro2A cells were seeded onto 9 mm borosilicate glass coverslips and transfected with miR-193-OFF, either alone or accompanied by ant-193 or Scr, following the procedures outlined above. Coverslips were placed in a Nikon Eclipse Ti-U microscope chamber equipped with Plan Fluor l0x PhlDL and S Plan Fluor ELWD 40x DIC N1 objectives (Nikon). Coverslips were immersed in a static extracellular solution comprising (in mM): 140 NaCI, 4 KCI, 1.8 CaCl_2_, 2 MgCL_2_, and 10 HEPES, pH adjusted to 7.34-7.37 with NaOH.; osmolality 293 mmol/kg.

∼5 MΩ borosilicate glass micropipettes were filled with an intracellular solution consisting of (in mM): 140 KCI, 1 MgCI_2_, 0.5 CaCL_2_, 10 HEPES, pH adjusted to 7.3 with KOH, osmolality 309 mmol/kg. V-gated potassium channel currents were recorded using whole-cell voltage-clamp configuration, monitoring cell capacitance and series resistance. Recordings with series resistance >20 MΩ were discarded. Series resistance compensation was not applied. Cells were held at a resting potential of -80 mV for 50 ms, followed by a 200 ms voltage step delivered in 10 mV increments up to +20 mV. Each current step was followed by a 40 ms hyperpolarizing step to -100 mV before returning to baseline for an additional 40 ms. Signals were amplified using an Axopatch 200B amplifier (Molecular Devices) and Axopatch 203BU headstage, lowpass filtered at 2 kHz, and sampled at 10 kHz using Clampex Software (Version 10.7) and a Digidata 1440A (Molecular Devices). Prior to data acquisition, cells were allowed to calibrate for at least three minutes. Recordings were conducted at room temperature.

### Statistical analysis

All statistical analyses were conducted using either Python (Version 3.9) with the following packages and versions: pyABF (2.3.7), NumPy (1.25.1), pandas (2.0.3), stasmodels (0.14.0), Seaborn (0.12.2), Matplotlib (3.7.2), SciPy (1.11.1), skimage (0.21.0) (code for these analyses at https://github.com/Joschua21/miRsense OFF.git) or GraphPad Prism (v10).

Data were tested for normality using Shapiro-Wilk test and parametric or non-parametric tests selected accordingly. Singular outliers were identified using a Grubbs test. Individual statistical tests used are listed in the figure legends for each p value presented. Data are presented as mean±SEM.

## Data availability statement

Raw data is available upon reasonable written request to the corresponding author.

## Acknowledgments

GM was funded by the Epilepsy Research Institute (F2102 Morris, 2501P Morris) and the Royal Society (RGS\R2\222326). SS and NT were funded by the UKRI Medical Research Council (MR/V034758/1).

## Author contributions

JG, NT, JB performed experiments and analysed data; SS and GM designed the study; JG and GM wrote the manuscript; All authors approved the final version of the manuscript.

